# Transcription-independent hold of the G1/S transition is exploited to cope with DNA replication stress

**DOI:** 10.1101/2022.11.10.515958

**Authors:** Yue Jin, Guoqing Lan, Jiaxin Zhang, Haoyuan Sun, Li Xin, Qinhong Cao, Chao Tang, Xiaojing Yang, Huiqiang Lou, Wenya Hou

**Affiliations:** Shenzhen University General Hospital, South China Hospital, Marshall Laboratory of Biomedical Engineering and Carson International Cancer Center, Guangdong Key Laboratory for Genome Stability & Disease Prevention, Shenzhen University School of Medicine, Shenzhen 518060; State Key Laboratory of Agrobiotechnology, College of Biological Sciences, China Agricultural University, Beijing 100193, China; State Key Laboratory of Agrobiotechnology, College of Biological Sciences, China Agricultural University, Beijing 100193, China; Center for Quantitative Biology and Peking-Tsinghua Center for Life Sciences, Academy for Advanced Interdisciplinary Studies, Peking University, Beijing 100871, China; School of Physics, Peking University, Beijing 100871, China

**Keywords:** genome instability, replication stress, DNA damage response, the cell cycle, tumor suppressor

## Abstract

RB1 (retinoblastoma) members control the G1/S commitment as transcriptional repressors in eukaryotic cells. Here we uncover that an extra copy of *RB1* equivalent (*WHI7* or *WHI5*) is sufficient to bypass the indispensability of the central genomic checkpoint kinases Mec1^ATR^-Rad53^CHK1^ in *Saccharomyces cerevisiae*. Mec1-Rad53 directly phosphorylate Whi7/5, antagonizing their nuclear export or protein turnover upon replication stress. Through in vitro reconstitution, we show that Whi7 C-terminus directly binds and hinders S-CDK-Cks1 from processively phosphorylating Sic1. By microfluidic single-cell real-time quantitative imaging, we demonstrate that both Whi7 and Whi5 are required to flatten the degradation curve of the major S-CDK inhibitor Sic1 in vivo. These findings reveal an eclipsed transcription-independent role of Whi7 homologs, which is highlighted by genome integrity checkpoints to hold the G1/S transition instantly as a rapid response to unforeseeable replication threats.

**Key points:**

1. Whi7 overexpression bypasses the essential function of Mec1 and Rad53 in a transcription-independent way.
2. Whi7 is stabilized by checkpoint-mediated phosphorylation.
3. Whi7 binds and hinders S-CDK-Cks1 from multi-phosphorylation of Sci1, thereby prolonging Sic1 degradation and G1/S transition.

## INTRODUCTION

The cell cycle needs to be dynamically regulated according to the environmental cues. Under normal conditions, the G1-to-S decision (Start) is a critical cell fate determinant point, either entering the S phase (i.e., DNA replication) or exiting the cell cycle (quiescence, senescence, or differentiation) (1). It is precisely controlled by Cln1/2/3-dependent kinases (G1-CDKs), transcription suppressors Whi5 and Whi7, two transcription factor (TF) complexes (SBF and MBF) and RNA pol II in *Saccharomyces cerevisiae* (2–6). Besides sharing a common subunit Swi6, SBF and MBF contain distinct DNA binding subunits Swi4 and Mbp1, respectively (7,8). Besides an SBF repressor like Whi5, Whi7 also kidnaps Cln3-Cdc28 in the endoplasmic reticulum (ER) till late G1 (3). After transportation to the nucleus, Cln3-Cdc28 initially phosphorylates and releases Whi5/7 from Swi6. Once transcription of the SBF-controlled genes including *CLN1* and *CLN2* is triggered, a positive feedback loop of G1-CDKs-Whi5/7-SBF/MBF is established to drive coherent G1/S transition (4,5,9). SBF preferentially drives genes in cell cycle timing and morphogenesis, whereas MBF primarily controls S-cyclin *CLB5, CLB6* and DNA metabolism genes (10,11). SBF is released from gene promoters in S phase via S-CDK-dependent phosphorylation (12), whereas MBF is inactivated by transcriptional repressor Nrm1 homologous to Whi5/7 via negative feedback (13). Besides SBF, S-CDKs also target Sic1 (<u>S</u>ubstrate <u>I</u>nhibitor of <u>C</u>dk) after being primed by G1-CDKs (14–16). The phosphates are added to multisite in a processive manner via docking by phospho-adaptor Cks1 in the S-CDK complex. Such a processive phosphorylation process triggers the degradation of Sic1, the major 4 inhibitor as well as a substrate of S-CDKs, further amplifying the S-CDK activity via positive feedback (17–19). In mammals, hyper-activation of CDK4/6 or *RB1* mutations often abrogates the G1/S restriction point and leads to over proliferation and tumor predisposition (20–23). Therefore, CDK4/6 kinase inhibitors become the first clinically approved anti-cancer drugs targeting the G1/S transition (23–25).

Under perturbed conditions, the G1/S transition is also modulated by genome integrity checkpoints comprising DNA damage checkpoint (DDC) and DNA replication checkpoint (DRC) (26–28). DDC acts against DNA damage throughout the cell cycle, whereas DRC functions against replication stress exclusively during S phase when genomic DNA is exposed and becomes most vulnerable (29,30). Partially intersecting, DRC and DDC consist of the Mec1^ATR^-Rad53^CHK1^ and Tel1^ATM^-Chk1^CHK2^ kinase cascades, respectively. In *S. cerevisiae*, Mec1-Rad53 represents the major genome integrity checkpoint and their encoding genes are essential for cell viability (31). Replication stress (i.e., slow DNA synthesis) causes uncoupling between DNA unwinding and synthesis, which results in the accumulation of single-strand DNA (ssDNA) coated by RPA (29). Once recruited and activated by RPA-ssDNA, the Mec1-Rad53 kinase cascade prevents cell cycle progression and rewires gene transcription to deal with replication stress (30).

Both DNA damage and replication stress inhibit the cell cycle progression by reducing CDK activities largely at the transcriptional level. In the presence of the RNR inhibitor hydroxyurea (HU), on the one hand, Rad53 phosphorylates Swi6 and Swi4 to down-regulate SBF target genes, including G1 cyclin genes *CLN1* and *CLN2*, to inhibit the G1/S transition in *S. cerevisiae* (32–35). Double deletion of *CLN1* and *CLN2* could cause DNA replication delays in a Sic1-dependent manner, thus bypassing the lethality of *mec1*Δ (36), reminiscent of the importance of the G1/S delay to cope with replication defects. On the other hand, Rad53 can phosphorylate Nrm1 and release it from Swi6 so that the MBF target genes can be continuously expressed in S phase (10,11). By regulating SBF and MBF in such an opposite manner, cells gain a golden time window to re-express the DNA replication/repair-related genes and allow these proteins to fix the problems under replication threats. This mechanism is conserved in human cells; CHK1 inhibits E2F6 repressor function to maintain the E2F-dependent transcription of the G1/S genes (37). E2F6 can determine the maximal amount of DNA a cell can synthesize per unit time (replication capacity) during S phase in a transcription-dependent manner (38).

In the current study, through a cross-species genetic screen, we have identified human *RB1* and budding yeast equivalents *WHI7* and *WHI5* as evolutionarily conserved dosage suppressors of the *mec1* or *rad53* mutants. Unanticipatedly, we found that Whi7 and Whi5 function in a transcription-independent manner. Whi7 and Whi5 are phosphorylated by the Mec1-Rad53 checkpoint, which antagonizes their nuclear export or protein turnover. Mechanistically, through in vitro reconstitution, we showed that Whi7 can directly bind Cks1, the S-CDK processivity factor, and thereby hinder S-CDK-Cks1 to processively phosphorylate Sic1, the major S-CDK inhibitor. Using microfluidic fluorescent live-cell microscopy, we showed that Whi7/5 redundantly flatten the Sic1 degradation under stress. These findings uncover a long-hidden mechanism by which G1/S transcriptional repressors are exploited as S-CDK-Cks1 inhibitors, thus prolonging the G1/S transition to deal with replication stress.

## RESULTS

### G1/S transcriptional repressors partially replace the essential roles of genome integrity checkpoint kinases

Since most of the known suppressors of *mec1* or *rad53* mutants in *Saccharomyces cerevisiae* are not well conserved in higher eukaryotes, we designed a cross-species dosage suppressor screen using a human cDNA library to identify the evolutionarily conserved targets of genome integrity checkpoints. In order to enrich putative checkpoint effectors, we prepared total mRNA from HEK293T cells 1 hour after 5 mM HU treatment. cDNA was then obtained by reverse transcription and cloned into a yeast expression plasmid. To minimize the possible toxic effect, we applied the *RNR3* promoter (*RNR3pr*) which is primarily induced by genotoxins. Although deletion of *SML1*, encoding an Rnr1 inhibitor, suppresses the lethality of *mec1*Δ (42), the *mec1*Δ*sml1*Δ cells barely grew in the presence of very low doses of HU (6 mM) (Figure 1A). However, wild-type (WT) budding yeast cells are usually resistant to over 200 mM HU. We then selected the plasmids that enabled the *mec1*Δ*sml1*Δ cells to grow on a plate containing 6 mM HU. The human *RB1* gene, encoding retinoblastoma tumor suppressor RB1, was identified as a putative dosage suppressor in a pilot screen (Figure 1A).

**Figure 1.**
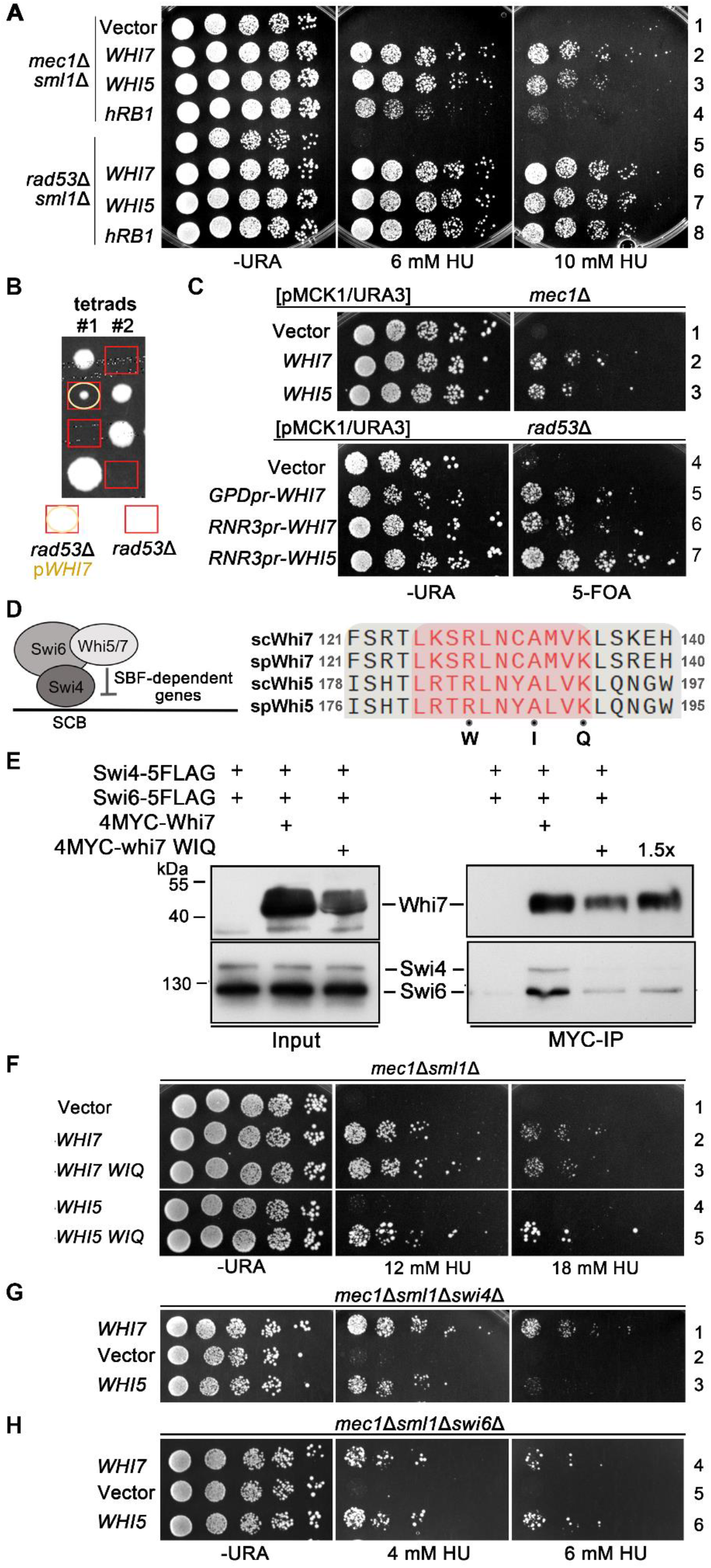
RB1 family members rescue *mec1*Δ or *rad53*Δ in a transcription-independent manner. (A) Overexpression of *WHI7, WHI5* or human *RB1* through the *RNR3* promoter suppresses the HU sensitivity of *mec1*Δ*sml1*Δ or *rad53*Δ*sml1* Δ. 5-fold serial dilutions of yeast cells containing the indicated plasmid were spotted onto the plates supplemented with different concentrations of HU. Plates were incubated at 30°C for 48 h before photography. (B) Overexpression of *WHI7* bypasses the essential role of *RAD53*. The plasmid expressing *WHI7* was transformed into *RAD53*^+/-^ heterozygotic diploid yeast strain, followed by tetrad dissection and genotype analysis. (C) An extra copy of either *WHI7* or *WHI5* rescued the lethality of both *mec1*Δ and *rad53*Δ. A single-copy plasmid pRS313 expressing *WHI7* or *WHI5* under the endogenous or *RNR3* promoter was transformed into the *mec1*Δ or *rad53*Δ haploid strain containing the *pRS426-MCK1* plasmid. Cells were spotted on SC-URA-LEU and SC-LEU+5-FOA plates for 5-fold serial dilution experiments. Photographs were taken after 72 h of culture in a 30°C incubator. (D) Whi7 and Whi5 transcription repressors restrict G1/S gene expression via the conserved GTB motif (in red). In their WIQ mutants, three invariable RAK residues are substituted by WIQ. (E) Whi7 interacts with Swi6 and Swi4 through the GTB motif as well. The pRS316-Whi7, pRS316-4MYC-Whi7 and pRS316-4MYC-whi7 WIQ plasmids were transformed into Swi6-5FLAG Swi4-5FLAG *whi7*Δ background strain. Immunoprecipitation was performed using MYC-Nanoab-Agarose. After three washes with high-salt (300 mM NaCl) buffer, the bound proteins were separated by 12% SDS-PAGE before immunoblotting. (F) Transcription-defective mutations render Whi5 a more potent checkpoint activity. WT or mutant *WHI7/WHI5* was overexpressed in *mec1*Δ*sml1*Δ. WIQ indicates mutations of the GTB motif as shown in (C). 2-fold serial dilutions were performed as described in (A). Before photography, plates were incubated at 30°C for 24 h in the absence of HU and 72 h in the presence of HU, respectively. (G, H) The checkpoint function of Whi7 and Whi5 is independent of TF Swi6 or Swi4. *WHI7* or *WHI5* was overexpressed in *mec1*Δ*sml1*Δ*swi4*Δ (G) or *mec1*Δ*sml1*Δ*swi6*Δ (H). 5-fold serial dilution assays were performed as described in (A).

Budding yeast has two functional *RB1* equivalents, *WHI5* and *WHI7*. We then introduced *WHI5* or *WHI7* under *RNR3pr* and also noticed suppression on the HU sensitivity of *mec1*Δ*sml1*Δ (Figure 1A; Supplementary Figure S1A, red square curve). Meanwhile, *RNR3pr*-driven *RB1, WHI5* or *WHI7* displayed a similar effect on *rad53*Δ*sml1*Δ as well (Figure 1A). Under constitutive overexpression driven by the *GPD* promoter (*GPDpr*), *WHI7*, particularly *WHI5*, was toxic under normal conditions (Supplementary Figure S1B, left panel), consistent with their canonical cell cycle inhibitory roles. In stark contrast, both of them dramatically increased cell viability in the presence of HU (Supplementary Figure S1B, right panel). These results indicate that Whi7 and Whi5 enhance cell growth under perturbed conditions whereas they restrict cell growth under normal conditions. In order to test the effect without introducing *SML1* deletion, we performed the tetrad dissection assays of the *rad53*Δ/*RAD53* diploid cells harboring a *WHI7* plasmid. After sporulation, four spores were separated under a microscope before genotyping. Among all *rad53*Δ spores, only those carrying the *WHI7* expression plasmid became viable (Figure 1B). Similarly, *WHI7* has been isolated as a weak <u>s</u>uppressor of *<u>r</u>ad53* <u>l</u>ethality (also named *SRL3*) in a previous large-scale genetic screen, whereas its mechanism remains unaddressed (44). To further examine whether *WHI7* or *WHI5* can compensate for the lethality of *mec1*Δ or *rad53*Δ, we introduced the *WHI7* or *WHI5* plasmid into these cells, whose viability can be supported by an *MCK1* plasmid as reported previously (43). After removal of the *MCK1::URA3* plasmid by 5-fluoroorotic acid (5FOA), we noticed that either *WHI7* or *WHI5* plasmid is sufficient to support the growth of both *mec1*Δ and *rad53*Δ cells (Figure 1C). Together, these data allow us to conclude that an extra copy of *WHI7* or *WHI5* is sufficient to bypass the essential role of both *MEC1* and *RAD53*.

### The checkpoint function of Whi7 and Whi5 does not depend on G1/S transcription

Given that Whi7 and Whi5 are transcriptional suppressors; we would like to know whether their checkpoint function also depends on the transcriptional activity. To this end, we mutated the conservative <u>G</u>1/S <u>t</u>ranscription <u>f</u>actor <u>b</u>inding (GTB) motifs (R, A, K residues) to WIQ (Figure 1D), which is known to abrogate the interaction of Whi5 with Swi6 (45). As shown by the coimmunoprecipitation experiments, WIQ mutations also curbed the association of Whi7 with Swi6 and Swi4 TFs (Figure 1E), indicating that Whi7 also adopts the GTB motif to regulate SBF-dependent transcription. Strikingly, *whi7-WIQ* retained full suppression capability (Figure 1F; Supplementary Figure S1A, cyan triangle curve; S1C row 6), whereas *whi5-WIQ* exhibited an apparent stronger suppression than wild-type (WT) (Figure 1F rows 4-5). These results suggest that the transcriptional repression activity is dispensable (if not deleterious) for their checkpoint function. Similarly, when the major SBF TF complex component Swi6 or Swi4 was omitted, *WHI7* or *WHI5* overexpression preserved the suppression capability (Figure 1G, H and Supplementary Figure S1D, E), supporting that G1/S transcription might be not required for their checkpoint function either. These data corroborate that Whi7 and Whi5 have a genome checkpoint function through a transcription-independent manner unanticipatedly.

### Whi7 is a direct target of genome integrity checkpoint

Because Whi7 shows a relatively potent checkpoint activity with less cytotoxicity, we next took Whi7 as an example to address the checkpoint function of G1/S transcriptional repressors in yeast. Whi7, a serine and threonine-rich protein (Figure 2A), is hyperphosphorylated by G1-CDKs to relieve its transcriptional repression during normal G1/S transition (3,4,46). To address whether Whi7 phosphorylation is modulated by replication stress, we probed the endogenous Whi7 protein carrying a 5FLAG tag by immunoblotting. Post HU treatment, the hyperphosphorylated Whi7 species increased substantially (Supplementary Figure S2A, lanes 1-2). Because HU treatment inhibits *Cln1/2* expression (33), HU-induced phosphorylation of Whi7 is unlikely due to CDKs. To further eliminate the interference of CDK-mediated phosphorylation, we mutated all putative CDK-sites (13 S/TP sites) in Whi7 to alanine. In the absence of 200 mM HU, hyperphosphorylation disappeared in whi7-13AP compared to WT Whi7 (Figure 2B, compare lane 2 to 1). After HU treatment, relatively slow-migrating whi7-13AP appeared (Figure 2B, lanes 3-4; Supplementary Figure S2A, lanes 3-9). However, slow-migrating whi7-13AP as well as the overall Whi7-13AP protein levels reduced in *mec1*Δ*sml1*Δ or *rad53* Δ*sml1*Δ cells (Figure 2B, compare lanes 5-10 with 2-4). After a better separation with a high-resolution Phos-tag gel shown in Figure 2C, Whi7 underwent multisite phosphorylation largely in a Rad53-dependent manner after HU treatment (lanes 1-4). Of note, despite being catalyzed by distinct kinases, the number of phosphorylated sites seemed not significantly changed with or without HU. These results indicate that Whi7 is phosphorylated at multiple sites in a checkpoint-dependent manner when cells encounter replication stress.

**Figure 2.**
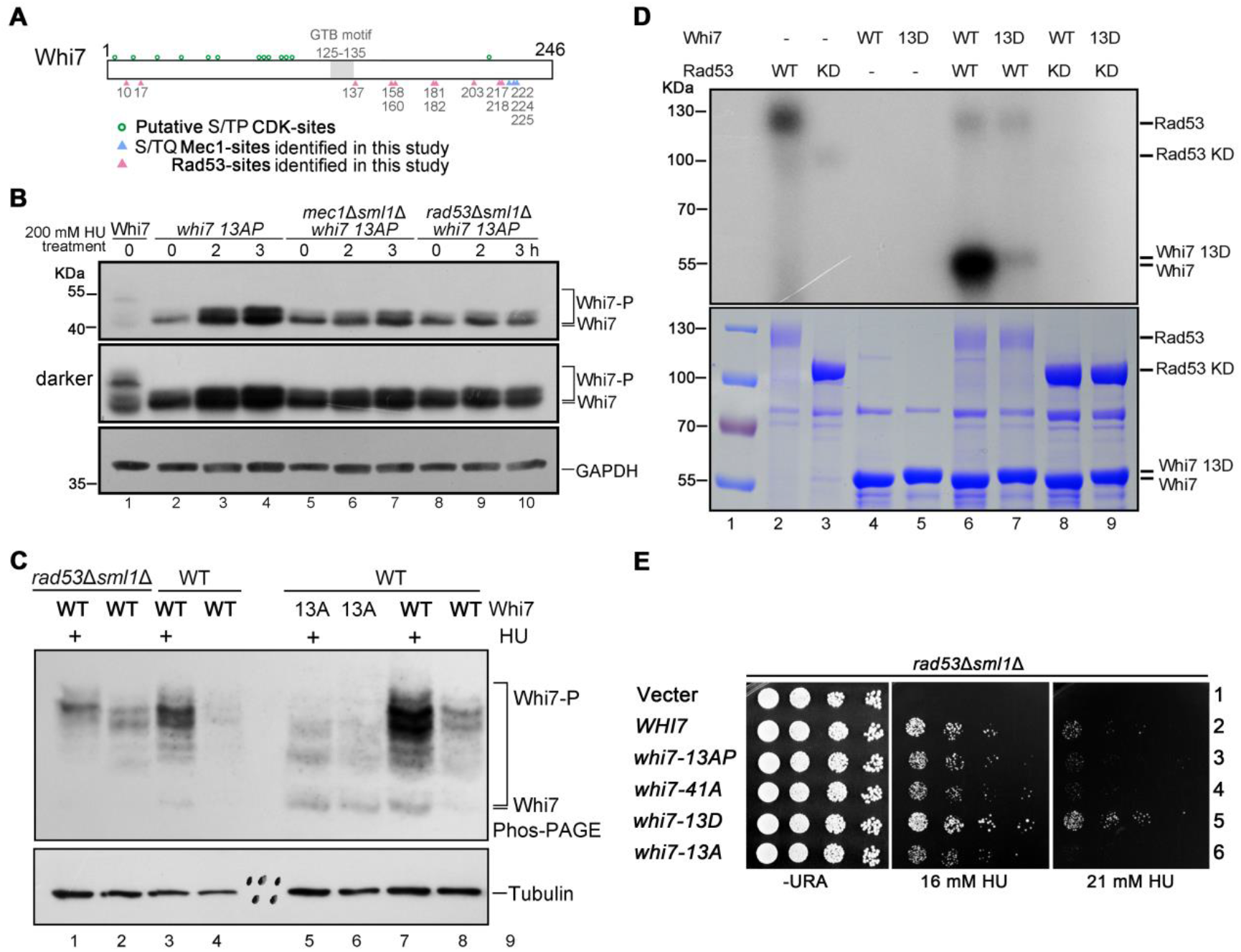
Whi7 is a direct target of Mec1 and Rad53 kinases. (A) Whi7 integrates multiple potential kinase signals other than CDKs. (B) Phosphorylated and total Whi7 are induced by HU largely via Mec1-Rad53. WT, *mec1*Δ *sml1*Δ or *rad53*Δ *sml1*Δ cells were collected after 200 mM HU treatment for the indicated time. The endogenous Whi7 carrying a 5FLAG tag in these cells was detected by immunoblots using an anti-FLAG antibody. Whi7-13AP represents that a total of 13 potential CDK sites (S/TP) of Whi7 are substituted by alanine. GAPDH serves as a loading control. (C) The main checkpoint-dependent phosphorylation sites of Whi7. The endogenous Whi7 or mutant whi7-13A (non-phosphorylatable by Mec1 and Rad53) was tagged by a 5FLAG tag. Cell lysates were separated by SDS-PAGE in the presence or absence of Phos-Tag before immunoblotting. Tubulin serves as a loading control. (D) Whi7 is directly phosphorylated by Rad53 *in vitro*. Recombinant His6-Rad53, His6-rad53-KD (K227A), GST-Whi7 and GST-Whi7-13D were expressed and purified from *E. coli*. Kinases and substrates were incubated in the presence of γ-^32^P-ATP as detailed in Methods. The samples were subjected to autoradiography after being resolved in an 8% polyacrylamide gel with SDS. The Coomassie Brilliant Blue (CBB)-stained gel shows the proteins in each reaction. (E) Phosphorylation of Whi7 facilitates its checkpoint functions. Dosage suppression assays were basically conducted as described in Figure 1A, except that various *whi7* mutants were applied. *whi7*-13AP indicates alanine substitutions of all putative CDK sites, whereas 41A represents alanine substitutions of all serine and threonine residues except 13 CDK sites. 13D and 13A indicate mutations of 13 serine and threonine residues (T10, S17, S137, S158, S160, S181, S182, S203, S217, S218, T222, S224, T225) of putative Rad53 or Mec1 sites to aspartic acid and alanine, respectively. 5-fold serial diluted samples were grown at 30°C for 72 h before photography.

We noticed that Whi7 harbors a ‘TQSTQ’ (a.a. 222-226) motif in its C-terminus (Whi7C, a.a. 217-246) (Supplementary Figure S2B). We next tested whether Whi7 is a substrate of Mec1 through in vitro kinase assays using purified proteins in the presence of γ–^32^P-ATP. As evidenced by autoradiography, a large amount of ^32^P was incorporated into recombinant Whi7 protein after incubation with Mec1 (Supplementary Figure S2C, lane 6). When three S/T residues within ‘TQSTQ’ were substituted by aspartic acid, whi7-3DQ showed a significantly reduced phosphorylation (Supplementary Figure S2C, lane 7). These results indicate that these S/TQ sites within Whi7C represent the main phosphorylation sites by Mec1. Functionally, overexpression of *whi7-3AQ* or *whi7* (1-221) displayed a slightly weaker suppression on HU sensitivity of *mec1*Δ*sml1*Δ and *rad53*Δ*sml1*Δ compared with that of *WHI7* WT or phospho-mimetic *whi7-3DQ* (Supplementary Figure S2D). These data suggest that Whi7 is directly phosphorylated by Mec1 for efficient replication stress alleviation.

Using a similar strategy, we showed that Whi7 is also directly phosphorylated by Rad53, but not rad53-KD (kinase-defective mutant) (Figure 2D, compare lanes 6 and 8). To identify Rad53-catalyzed phosphorylation sites, recombinant Whi7 was incubated with Rad53 in the presence of ATP prior to liquid chromatography-tandem mass spectrometry (LC-MS/MS). A total of 10 putative phosphorylation sites were detected (Supplementary Figure S2E). Since phosphorylation sites are often redundant, we next mutated all putative Rad53-sites to aspartic acid in the *whi7-3AQ* or *whi7-3DQ* background. Whi7-13A and Whi7-13D nearly lost phosphorylation in vivo (Figure 2C, lanes 5-8) and in vitro (Figure 2D, compare lane 6 with 7), respectively. Functionally, *whi7-13D* (mimicking phosphorylation by Mec1 and Rad53) acted as a moderate stronger dosage suppressor of the *rad53* mutant than *WHI7* WT (Figure 2E, compare row 5 to 2), whereas *whi7-13A* (non-phosphorylatable by Mec1 and Rad53) became a mild weaker suppressor (compare row 6 to 2). When we further mutated all S/T residues other than the S/TP sites throughout Whi7, *whi7-41A* displayed a similar suppression to *whi7-13A* (Figure 2E, compare rows 4 and 6), indicating that these 13 S/T residues of Whi7 represent nearly all redundant phosphorylation sites of the Mec1-Rad53 pathway. These data suggest that Whi7 is a direct substrate of Mec1-Rad53 in replication stress response.

### Checkpoint-mediated phosphorylation stabilizes Whi7

We next asked whether checkpoint-dependent phosphorylation of Whi7 affects its protein turnover like the CDK-mediated one. Because the protein levels of Whi7 fluctuate during the cell cycle, we first used the α-factor to arrest the yeast cells in G1 phase. Then cells were released into fresh media supplemented with cycloheximide (CHX) to inhibit de novo protein synthesis. Under normal conditions, Whi7 declined rapidly in about 30 min release from G1 (Figure 3A). In the presence of 200 mM HU, Whi7 declined at a significantly slower rate (Figure 3B), suggesting that replication stress may extend the half-life of Whi7 protein. To exclude the interference of CDK-triggered Whi7 degradation during the normal G1/S transition, we used a *whi7-13AP* mutant strain described in Figure 2B. As expected, *whi7-13AP* mutant protein, no longer a CDK substrate, was degraded much slower than Whi7 WT (Supplementary Figure S3A, compare lanes 6-10 with 1-5). Moreover, Whi7-13AP remained relatively stable for more than 60 min after HU treatment (Figure 3C, lanes 1-4; 3D, red dot curve). However, when combined with whi7-3AQ, Whi7-13AP-3AQ was no longer stabilized even in the presence of 200 mM HU (Figure 3C, lanes 5-8; 3D, cyan square curve). These data indicate that Whi7 is stabilized by genome integrity checkpoint under stress.

**Figure 3.**
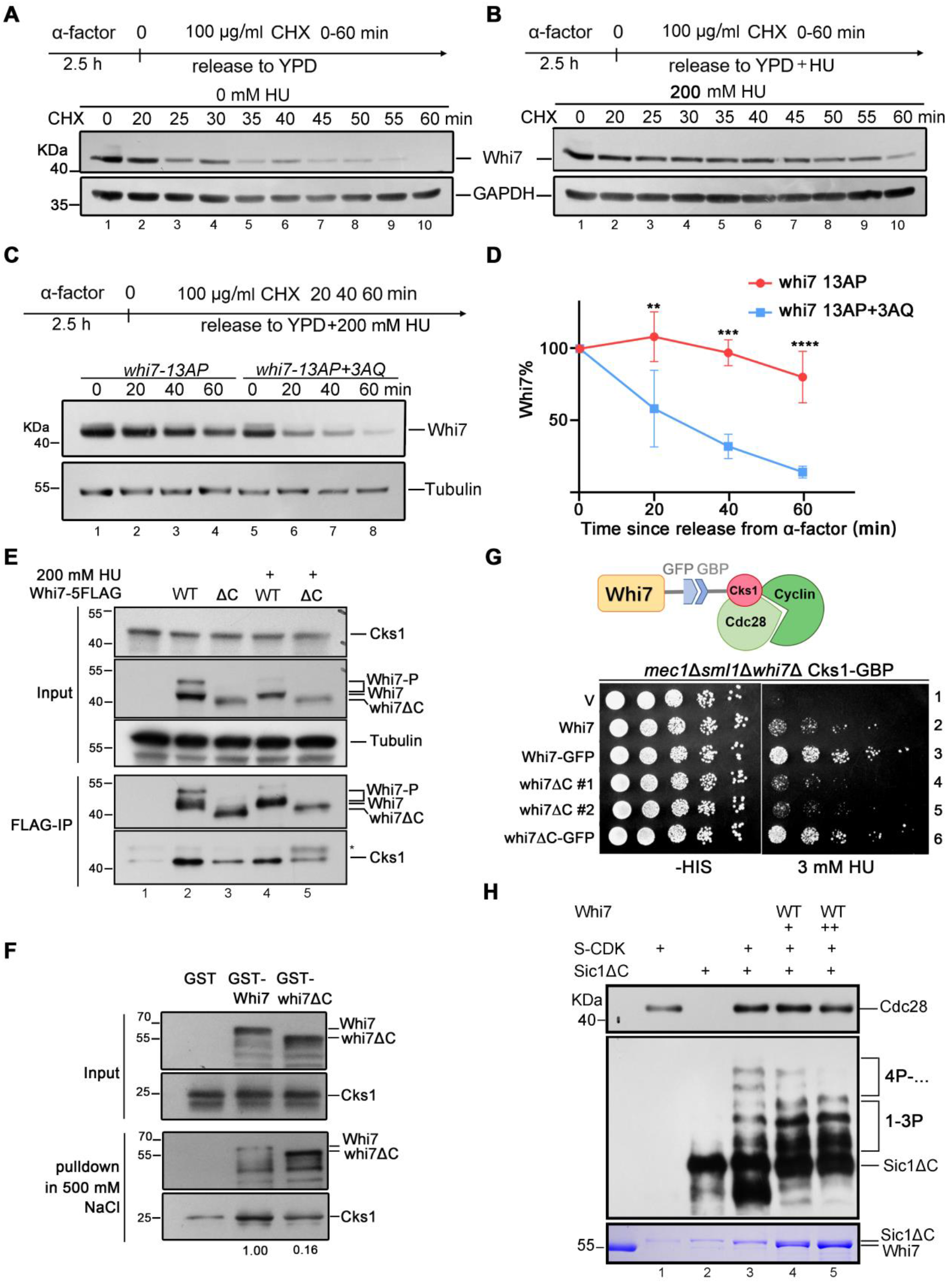
Checkpoint-mediated Whi7 phosphorylation counteracts protein turnover. (A, B) Whi7 is stabilized in a checkpoint-dependent way in response to HU. Cells carrying the endogenously tagged whi7-13AP or whi7-13AP-3AQ were arrested in G1 by α-factor and released into fresh media containing 200 mM HU. 100 μg/ml of cycloheximide (CHX) was then added at t=0; samples were collected every 20 mins. Cell lysates were analyzed by immunoblotting. Tubulin serves as a loading control. The relative amounts of Whi7 compared with Tubulin were quantified by BioRad Quantity-One. (C, D) Checkpoint-mediated phosphorylation stabilizes Whi7 in the presence of HU. The half-lives of *whi7-13AP* and *whi7-13AP+3AQ* were quantified from three independent repeats as described above. The statistical significance was calculated via two-way ANOVA analysis, **, p < 0.01; ***, p < 0.001; ****, p < 0.0001. (E) Whi7 interacts with Cks1 primarily via its C-terminus. Immunoprecipitation was conducted as described in Figure 1E. (F) Whi7 directly associates Cks1 through its C-terminus. Cks1-ALFA was purified from yeast cells and mixed with recombinant Whi7 in glutathione beads. Before elution, non-specifically bound proteins were washed with a high-salt buffer containing 500 mM NaCl. The bound proteins were analyzed via SDS-PAGE and immunoblotting. (G) Whi7C deletion compromises the dosage suppression effects, which can be restored entirely via direct Whi7-Cks1 tethering. (H) Whi7 directly inhibits Sic1 hyperphosphorylation by S-CDKs. S-CDK complexes were purified from S-phase yeast cells. Recombinant GST-Whi7 was expressed and purified from *E. coli* BL21. The products of the in vitro kinase reactions were subjected to 10% SDS-PAGE and Phos-tag PAGE followed by immunoblotting (two upper panels) and Coomassie Brilliant Blue staining (bottom panel), respectively.

### Whi7 binds and inhibits S-CDKs-Cks1 to phosphorylate Sic1 processively

What’s the role of the stabilized Whi7? Interestingly, Whi7 was reported to be co-purified with Cks1 in a previous large-scale protein complex screen study (52). We validated their interaction by coimmunoprecipitation at their endogenous protein levels (Figure 3E). Notably, Cks1 coprecipitated efficiently with Whi7 regardless of the presence or absence of HU, implying that replication stress enhances protein stability of Whi7 but not Whi7-Cks1 interaction. Furthermore, their association was direct and stable up to 500 mM NaCl as evidenced by in vitro pulldown assays using purified recombinant proteins (Figure 3F). Through a series of truncation screens, we found that the loss of Whi7C (a.a. 217-246) significantly compromised its interaction with Cks1 as shown by both IP (Figure 3E, lanes 3 and 5) and in vitro pulldown experiments (Figure 3F). Functionally, Whi7C was necessary for dosage suppression and bypassing *RAD53* (Figure S1C, row 5; Supplementary Figure S3B, compare rows 4 and 2). These data suggest that Whi7C-mediated association with Cks1 is involved in replication stress response.

If the compromised interaction with Cks1 is a genuine cause of the checkpoint defect in whi7ΔC, we could restore its function by reinforcing their interaction. To test this notion, we tagged the endogenous Whi7 with GFP and Cks1 with GBP (GFP binding protein), respectively (Figure 3G). Unexpectedly, an extra copy of *WHI7* was no longer necessary; the endogenous Whi7 was sufficient to suppress the HU sensitivity of *mec1*Δ*sml1*Δ as long as it carried a GFP to bind Cks1-GBP (Figure 3G, compare row 3 to 2). Importantly, such a direct tethering also fully restored the checkpoint capability of whi7ΔC to that of WT levels (Figure 3G, rows 6 and 3). Our results indicate that the checkpoint function of Whi7 is attributed to its association with Cks1.

Cks1 is the processivity subunit of the S-CDK complex required for hyperphosphorylation of the substrates like Sic1, the primary S-CDK inhibitor during G1/S transition (16). Therefore, we next examined whether this activity is modulated by Whi7 by in vitro reconstitution of Sic1 phosphorylation catalyzed by CDKs. Native CDK proteins were purified from the early S-phase yeast cells overexpressing the S-phase cyclin Clb5 (Supplementary Figure S3C). A truncated version of recombinant Sic1 (sic1ΔC) that escapes from degradation was purified and incubated with CDKs. A Phos-tag gel was used to separate multisite phosphorylation with a better resolution. Various phosphorylated sic1ΔC species were clearly observed (Figure 3H, compare lane 3 to 2), confirming a successful reconstitution of the reaction in vitro. Intriguingly, the addition of purified Whi7 or Whi7-13D proteins substantially reduced Sic1 hyperphosphorylation in a dose-dependent manner (compare lanes 3-5), correlating well with the pattern when the phospho-adaptor Cks1 within S-CDKs is inactivated in a previous study (16). An alternative explanation could be due to the competitive inhibition by Whi7 as the substrate of CDKs, which are unlikely due to at least three reasons. First, Whi7 homologs are the substrate of G1-CDKs but not S-CDKs (2). Moreover, they do not inhibit G1-CDKs as the substrates either. Second, as shown in Figure 3H, it is worthy to note that Whi7 specifically inhibits hyperphosphorylation (more than 4P, Cks1-dependent), but not hypophosphorylation (e.g. fast prime phosphorylation by G1-CDKs). Thus, these data suggest that Whi7 can directly inhibit the processive Sic1 phosphorylation catalyzed by S-CDK-Cks1.

### Whi5 is retained in the nucleus by checkpoint

Since Whi5, the paralog of Whi7, also acted as a dosage suppressor of the *mec1* or *rad53* mutants in a transcription-independent way in the presence of HU (Figure 1). We reasoned that they might share a redundant role in genome integrity checkpoint. In contrast to Whi7, Whi5 is highly stable and primarily regulated via nucleoplasmic shuffling (5,6). Therefore, we next examined whether its nuclear localization is regulated by genome integrity checkpoint. To this end, Whi5-GFP is labeled at its genomic locus and analyzed by a live-cell microscope equipped with microfluidic devices. Yeast cells were loaded into different microfluidic devices fed with a constant flow of the liquid media. Fluorescent images were acquired at 3-minute intervals. Consistent with a previous report (15), Whi5 diffused from the nucleus into the cytoplasm within 12 min during normal G1/S transition (Figure 4A, upper panel; 4B, cyan dot curve). However, such diffusion was significantly delayed after the switch to media containing 200 mM HU (Figure 4A, lower panel; 4B red square curve). Strikingly, such a HU-induced delay disappeared in the absence of Rad53 (Figure 4C, lower panel; 4D), indicating the Rad53-dependent nuclear detention of Whi5. Given that Whi5 does not contain any S/TQ motifs; it is unlikely recognized by Mec1. However, Whi5, like its paralog Whi7, was indeed a direct substrate of Rad53 (Figure 4E, lane 3). These data suggest that the nuclear export of Whi5 is inhibited in a Rad53-dependent way. Similar to its paralog Whi7, Whi5 was capable to inhibit Sic1 hyperphosphorylation by S-CDK-Cks1 (Figure 4F, lane 6). Together, these data suggest that both Whi5 and Whi7 can inhibit Sic1 multi-phosphorylation although they are retained by checkpoint at the subcellular localization and protein levels, respectively.

**Figure 4.**
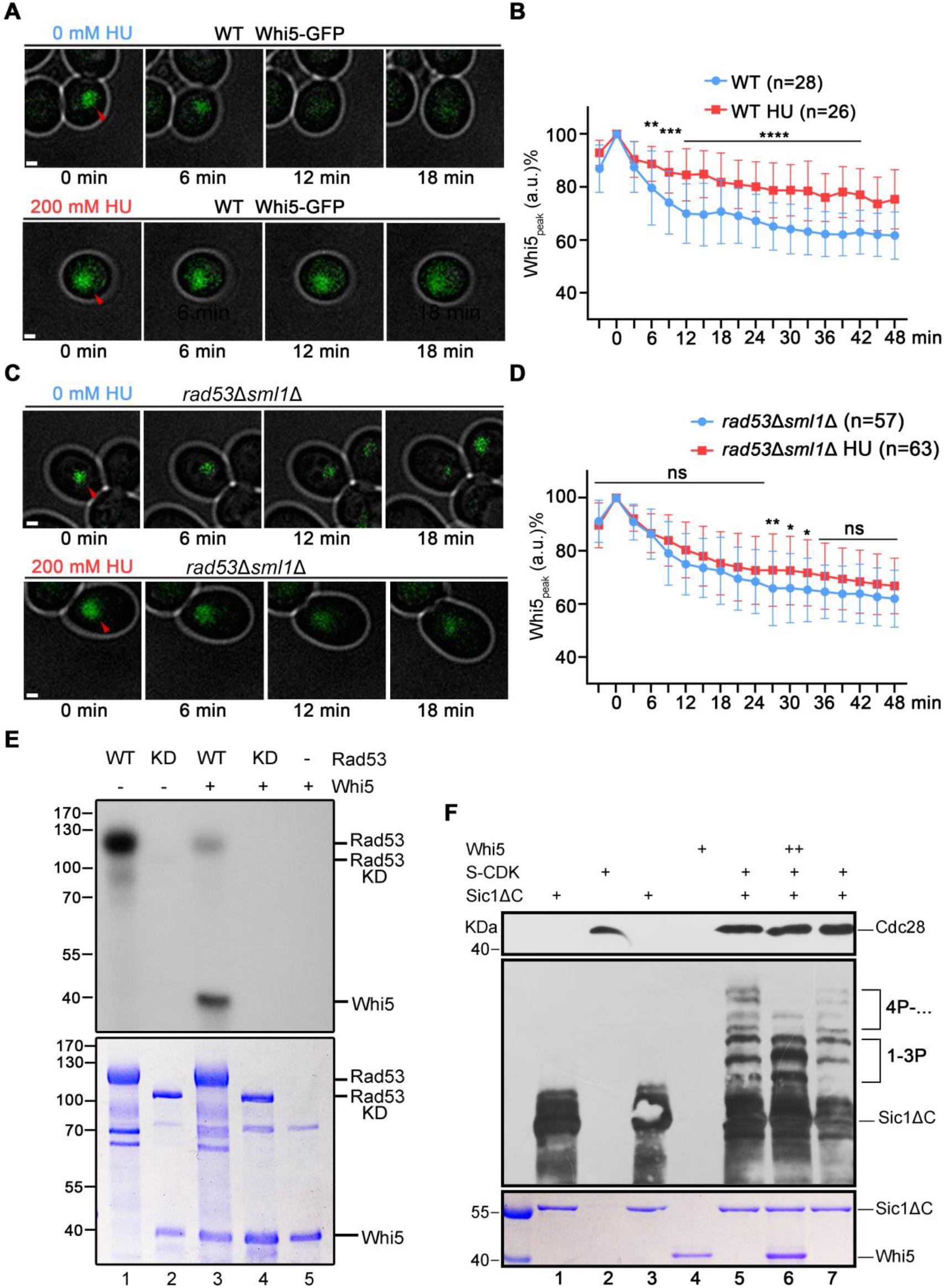
Whi7 and Whi5 inhibit the processive phosphorylation of Sic1 by S-CDKs-Cks1. (A-D) Whi5 is detained within the nucleus by Rad53. Yeast cells carrying an endogenous Whi5-GFP in WT (A, B) or *rad53*Δ *sml1*Δ (C, D) background were cultured in a microfluidic device supplemented with YPD medium with or without 200 mM HU at a constant flow rate. The mean fluorescence intensity of the brightest 5×5 pixel was quantified (B, D). The peak intensity of each cell was normalized to 100%. The statistical significance was calculated via two-way ANOVA analysis, *, p < 0.05; **, p < 0.01; ***, p < 0.001; ****, p < 0.0001. ns, no significant difference. (E) Rad53 phosphorylates Whi5 in vitro. In vitro kinase assays were performed as described in Fig 2D, using purified Whi5 and Rad53 kinase. (F) Whi5 directly inhibits Sic1 hyperphosphorylation by S-CDKs. S-CDK complexes were purified from S-phase yeast cells. Recombinant 6His-Whi5 was expressed and purified from *E. coli* BL21. The products of the in vitro kinase reactions were subjected to 10% SDS-PAGE and Phos-tag PAGE followed by immunoblotting (two upper panels) and Coomassie Brilliant Blue staining (bottom panel) as described in Figure 3H.

### Whi7 and Whi5 prolong the G1/S transition under stress

Multi-phosphorylation of Sic1 leads to its degradation (19,50). Therefore, we next this process is interrupted by Whi7 and Whi5 in vivo. We applied western blotting to measure the degradation of endogenous Sic1. The cells were synchronized in G1 by α-factor before the release into fresh media for the indicated time. In the absence of HU, Sic1 declined rapidly and disappeared within 40 min (Figure 5A, lanes 1-5; 5B, black curve). After 200 mM HU treatment, Sic1 remained for an additional 20 min and declined significantly slower (Figure 5A, lanes 6-9; 5B, red dot curve). Our results indicate that Sic1 degradation and G1/S transition are substantially delayed in response to HU. When both *WHI7* and *WHI5* were deleted, Sic1 was degraded at a similar rate to WT yeast cells under normal conditions (Figure 5A, lanes 10-14; 5B, grey diamond curve). However, in the presence of HU, Sic1 disappeared much faster in *whi5*Δ*whi7*Δ than in WT cells (Figure 5A, lanes 15-18; 5B, cyan square curve). Our data suggest that when cells encounter replication stress, Whi7 and Whi5 significantly slow down the Sic1 degradation whereas they only have a neglectable effect under normal conditions. To further validate this in a real-time way at the single-cell level, we quantified the endogenous Sic1-GFP levels through live-cell fluorescence imaging in a microfluidic device. This experiment bypassed the α-factor and CHX treatments and thus reflected a more physiological response under such borderline arrest situations. Analogously to the experiment described in Figure. 5A, Sic1 declined at nearly the same rate in WT and *whi5*Δ*whi7*Δ cells under normal conditions (Supplementary Figure S4A-C). After a switch to HU-containing media, Sic1 turnover was not affected in *whi5*Δ*whi7*Δ cells but was rapidly inhibited in WT cells (Figure 5C, 5D and Supplementary Figure S4C). Because the Sic1 levels reflect the G1/S transition (49), these data allow us to conclude that Whi7 and Whi5 postpone the Sic1 degradation and thereby G1/S transition in response to replication stress.

**Figure 5.**
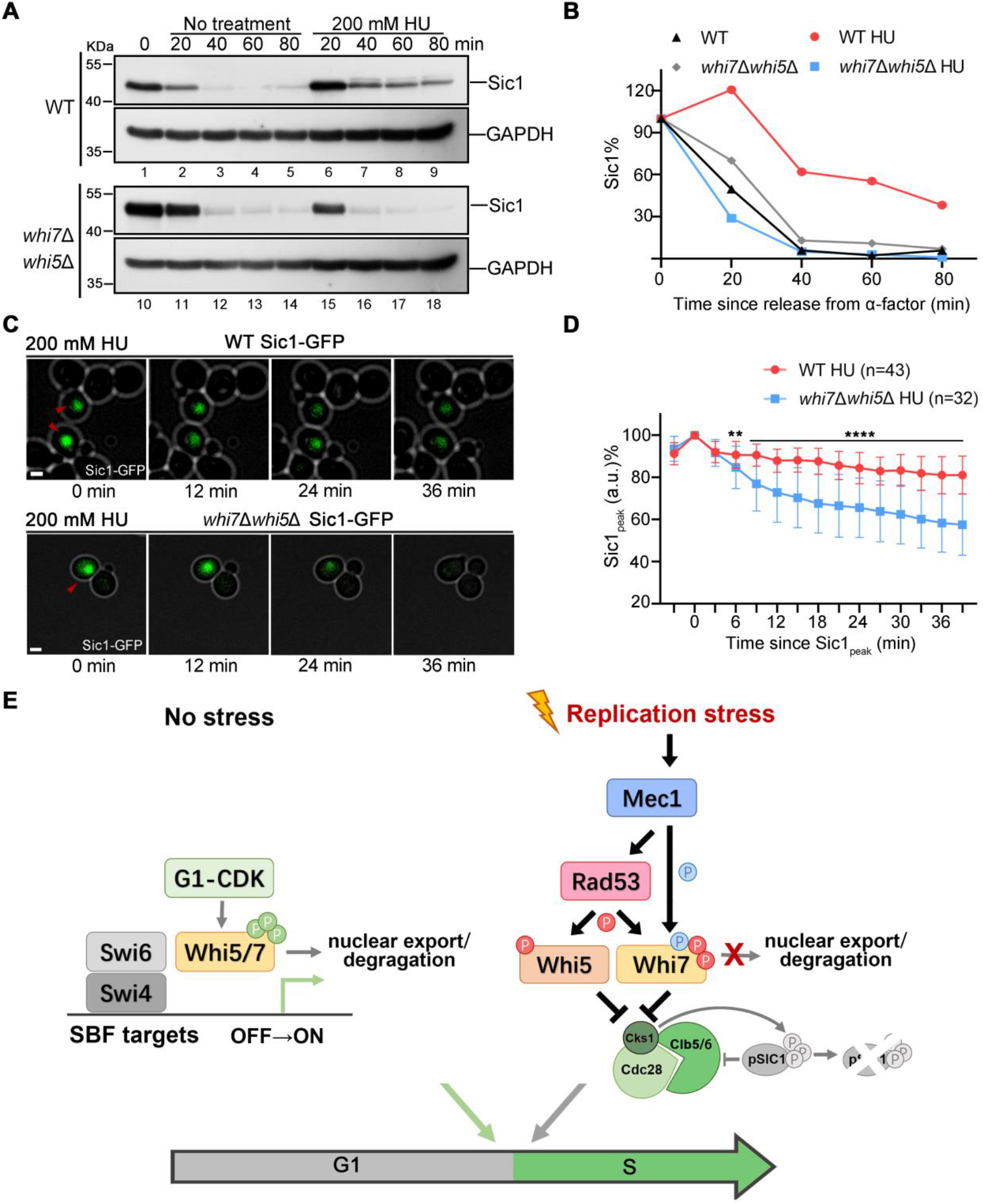
Whi7 and Whi5 inhibit the processive phosphorylation of Sic1 by S-CDK-Cks1. (A, B) Whi7/5 are required for HU-induced slower degradation of Sic1. The half-lives of Sic1 were measured as in Fig 3A. The Sic1 band density relative to the GAPDH loading was quantified and plotted against time after HU treatment (B). (C, D) Single-cell live-cell imaging of Sic1-GFP yeast cells after HU treatment. The averages of the Sic1-GFP signals from more than 20 cells are plotted against time since Sic1_peak_ (D). The statistical significance was calculated via two-way ANOVA analysis, **, p < 0.01; ****, p < 0.0001. (E) The dual functions of Whi5/7 in dynamic G1/S regulation. Under normal conditions, Whi5/7 primarily acts as transcriptional repressors. G1-CDKs phosphorylate Whi5/7, triggering their nuclear export or degradation, releasing the SBF/MBF-driven transcription and allowing the G1/S transition. Under replication stress conditions, Whi5/7 primarily functions as S-CDK-Cks1 processivity inhibitors. The checkpoint kinases Mec1-Rad53 phosphorylate Whi7 and Whi5 and kidnap them in the nucleus. Whi7 and Whi5 directly bind and inhibit S-CDKs from hyper-phosphorylating Sic1. This results in an instant G1/S delay as a fast response to replication stress.

Finally, we wanted to know whether the transcription-independent mechanism of Whi7 also applies under unperturbed conditions. Indeed, constitutive overexpression of *whi7*-WIQ significantly inhibited the G1/S transition even in the absence of stress (Figure S5). Such inhibition also required the Cks1-interacting motif, Whi7C, implying that the Cks1-inhibitory activity revealed in this study might also contribute to normal cell cycle control although it is eclipsed by its dominant transcription-inhibitory activity. In sum, in addition to transcription repression, Whi7 and Whi5 possess a neglected S-CDK-Cks1 inhibitory activity, which is exploited by genome integrity checkpoints to instantly postpone the Sic1 degradation and G1/S transition as a quick response to unforeseeable replication threats (Figure 5E).

## DISCUSSION

Whi7 and Whi5 are known to restrict the G1/S transition as transcriptional repressors. Here we report that they also modulate the G1/S transition through a transcription-independent mechanism under both perturbed and unperturbed conditions in budding yeast. The Mec1-Rad53 checkpoint detains Whi7 and Whi5, extending their service as inhibitors of Sic1 multi-phosphorylation by S-CDK-Cks1. By this means, the G1/S transition is prolonged to allow cells to deal with replication stress.

First, normal G1/S transition is stringently controlled by a belt-and-braces approach at transcription and post-translational levels. Because gene expression is notoriously leaky and stochastic in transcription and translation (53), the S-CDK inhibitory activities of Whi7 and Whi5 provide an extra valve to secure a proper cell cycle commitment.

Second, the division of labor between these two roles, S-CDK-Cks1 inhibition and transcriptional repression, can be rewired under different conditions. Under normal conditions, the S-CDK-Cks1 inhibitory activities of Whi5/7 are completely eclipsed by their transcriptional repression. However, under perturbed conditions, S-CDK-Cks1 inhibition rather than transcriptional repression is exploited by genome integrity checkpoints to prolong the G1/S transition. It explains why their transcription-independent role has remained buried for decades. Such a role transition may have some crucial physiological implications. For instance, the S-CDK-Cks1 inhibitory activities of Whi7 and Whi5 can exert a direct, rapid, cost-effective and potentially reversible action, whereas the transcription-dependent response is indirect, delayed, long-lasting and high-cost (54). A post-translational modification-triggered ready-made molecular brake might be of particular significance to surviving a threatening and everchanging stimulus. In this term, our findings may modify the current view of dynamic cell cycle regulation according to environmental cues.

Third, the Mec1-Rad53 pathway also maintains MBF-dependent transcription by targeting Nrm1 (10,11). It stimulates a class of MBF target genes, including both *RNR1* and S-cyclin genes *CLB5/6*. The latter ones need to be kept transiently inhibited, at least partially by Whi5 and Whi7, until the stress is alleviated.

Fourth, the division of labor between Whi7 and Whi5 is also changed under perturbed conditions. Albeit largely redundant, Whi5 plays a dominant role in sensing intracellular cues (i.e., cell volumes) and controlling the cell cycle under normal conditions, whereas Whi7 is relatively prone to finetuning the cell cycle according to various extracellular cues (55). Correspondingly, Whi5 protein is highly stable while Whi7 undergoes dynamic turnover (4,41).

Human RB1 likely has a conserved role as their yeast counterparts. Treatment with cisplatin, etoposide, or mitomycin C inhibits the S phase progression in RB1^+/+^ but not in RB1^-/-^ mouse embryo fibroblasts (56). RB1 is also directly targeted by ATM/ATR-CHK2/CHK1 (57). However, these studies have demonstrated that RB1 exerts another downstream effect, DNA repair, in a transcription-independent fashion (58,59). After being phosphorylated and loaded onto the DSB sites by the BRCT6 domain of TopBP1, the RB-E2F1 complex recruits several critical DNA repair factors such as BRG1 (a core ATPase subunit of the chromatin remodeler SWI/SNF (hBAF/PBAF)), MRN (MRE11-RAD50-NBS1) and histone acetyltransferases p300-CBP. Putting it all together, we propose that RB1 family proteins are responsible for at least two essential DDR effects, cell cycle inhibition and DNA repair (57,59). The S-CDK-Cks1 inhibitory function of RB1 members will extend our understanding of this tumor suppressor and related cancer etiology.

## MATERIAL & METHODS

### Strain construction

All yeast strains used in this study are isogenic with *S. cerevisiae* BY4741. Strain and plasmid information is summarized in Supplementary Table S1 and S2, respectively. All gene manipulation and protein tagging were made using Longtine’s vectors (39).

### Drug sensitivity assays

Log-phase growing cells (initial OD600 = 0.2) were spotted on YPD or synthetic media plates by five-fold serial dilution in the presence of the indicated concentrations of HU. Before photography, plates were incubated at 30°C for 48 h or 72 h.

### GST pulldown experiments

The *E. coli* lysates expressing the indicated proteins are mixed with glutathione-Sepharose beads in the presence of lysis buffer (50 mM Tris-HCl, pH7.5, 500 mM NaCl, 0.1 mM EDTA, 10% glycerol, 0.1% Triton X-100, 1 mM DTT, 1 mM PMSF, and protease inhibitors) for 1 h at 4°C. The glutathione agarose beads were washed extensively before the bound proteins were separated on 15% SDS-PAGE gels. Blots were probed with a monoclonal antibody against GST (1:3000) or ALFA (1:10000).

### Immunoprecipitation (IP)

Immunoprecipitation (IP) was carried out as described previously (40).

### In vitro kinase assays

His6-Rad53 and His6-rad53-KD (a kind gift from Dr. John Diffley) were purified using Ni-NTA chromatography (GE Healthcare). Recombinant GST-Whi7 and GST-Whi7 mutant proteins were purified using GST affinity chromatography. 3FLAG-Mec1 was precipitated from yeast cells by M2 beads (Sigma). The reaction was completed in 50 mM Tris, pH 7.5, 150 mM NaCl, 10 mM MgCl_2_, 5 μCi of γ-^32^P-ATP at 37°C for 30 min. Reactions were quenched by adding the SDS loading buffer and boiled for 10 min before SDS-PAGE and autoradiography.

CDK kinase assays were performed according to (16) with some modifications. In brief, GST-Sic1ΔC-ALFA was purified using GST affinity chromatography. S-CDK was obtained by Cdc28-5FLAG IP. We cultivated Cdc28-5FLAG yeast cells harboring *pRS425-Gal1-CLB5* plasmids in raffinose medium to logarithmic phase, then synchronized cells in G1 phase with α-factor. Cells were then released into the α-factor-free galactose medium to induce *CLB5* expression for 30 min to proceed into S phase. Proteins were detected by SDS-PAGE and Phos-Tag SDS-PAGE (8% SDS-PAGE supplemented with 40 μM Phos-Tag reagent), using anti-FLAG (Sigma) antibody and anti-ALFA (nano tag) antibody.

### Protein half-life assays

Cells were grown to log phase before adding 100 μg/ml (a dose allowing slow cell cycle progression) Cycloheximide CHX for the indicated time. Yeast cell extracts were prepared using the trichloroacetic acid (TCA) precipitation for SDS-PAGE and immunoblotting. The immunoblots with appropriate exposure were scanned and quantified using Quantity ONE (BioRad). The protein levels were normalized to the loading control.

### Use of a microfluidic device and time-lapse microscopy

Time-lapse microscopy mounted on the microfluidic device was carried out as described in (41). Briefly, yeast cells were loaded into the microfluidic device and fed into the culture medium at a constant rate of 66.7 μl/hour. Before imaging, cells were precultured in the microfluidic chip at 30°C for 2 h.

All images were captured by a Photometrics EMCCD Evolve512 at 3-minute intervals. Cell segmentation and fluorescent quantification were performed by Cellseg. We used the mean intensity of the brightest 5×5 to indicate the fluorescence intensity of Sic1-GFP at the moment. During one round of the cell cycle, the maximum fluorescence value was defined as Sic1_peak_ and normalized as 100%.

## Supplementary Data

Supplementary Tables S1–S2 and Figure S1-S5 are available online.

## ACKNOWLEDGMENTS

We’re grateful to Drs. John Diffley (The Francis Crick Institute), Hui Jiang (National Institute of Biological Sciences, Beijing) for yeast strains and plasmids, and the members of the Lou laboratory for discussion.

## Funding

This work was supported by the National Natural Science Foundation of China Grants (32161133015 and 32101039); National Key R&D Program of China (2019YFA0903900); Ministry of Science and Technology of China (2018YFA0900700, 2021YFF1200500); Natural science foundation of Guangdong province of China (2022A1515012495 and 2022A1515011208), Beijing Municipal Natural Science Foundation 5212010, SZU Top Ranking Project 86000000210 and Open Research Fund of the National Center for Protein Sciences at Peking University in Beijing.

## AUTHOR CONTRIBUTIONS

Conceptualization, H.L. and Y.J.; Methodology, X.Y, C.T.; Investigation, Y.J., J.Z., H.S., L.X. and W.H.; Resources, X.Y., W.H. and Q.C.; Writing-Original Draft, H.L. and Y.J.; Writing-Review & Editing, H.L., Y.J. and X.Y.; Visualization, Y.J., J.Z., H.L. and X.Y.; Supervision, H.L., C.T., X.Y. and Q.C.; Funding Acquisition, H.L., C.T., X.Y., Q.C., and W.H.

## Lead contact

Further information and requests for resources and reagents should be directed to and will be fulfilled by the Lead Contact, Huiqiang Lou.

## Declaration of interests

The authors declare no competing interests.

## Notes

### Competing Interest Statement

The authors have declared no competing interest.

