## Supplementary data for "Transcription-independent hold of the G1/S transition is exploited to cope with DNA replication stress"

Tables S1-S2, Figures S1-S5 and supplementary figure legends.

### Transcription-independent hold of the G1/S transition is exploited to cope with DNA replication stress

Yue Jin[1], Guoqing Lan[2], Jiaxin Zhang[1], Haoyuan Sun[3], Li Xin[3], Qinhong Cao[2], Chao Tang[3,4], Xiaojing Yang[3], Huiqiang Lou[1], Wenya Hou[1,*]

**Table S1. Yeast Strains used in this study**

| Lab name | Genotype |
|---|---|
| BY4741 | *MATa his3Δ1 LEU2Δ0 met15Δ0 URA3Δ0 lys2Δ0* |
| LXL190929025 | *BY4741 mec1Δ::KanMX sml1Δ::Leu2 hug1Δ::NatMX crt1Δ:: HygR* |
| Kim 07 | *BY4741 rad53Δ::KanMX sml1Δ::LEU2 whi7Δ::NatMx* |
| Kim 08 | *BY4741 mec1Δ::KanMX sml1Δ::LEU2 whi7Δ::NatMx* |
| Kim 12 | *BY4741 mec1Δ::KanMX sml1Δ::Leu2 rnr3Δ::HisMX* |
| Kim 29 | BY4741 *mec1Δ::KanMX sml1Δ::LEU2* |
| Kim 30 | *BY4741 rad53Δ::KanMX sml1Δ::LEU2* |
| Kim 106 | *BY4741 whi7-13AP-5FLAG::HisMX* |
| Kim 107 | *BY4741 rad53Δ::KanMX sml1Δ::LEU2 whi7-13AP-5FLAG::HisMX* |
| Kim 109 | *BY4741 mec1Δ::KanMX sml1Δ::LEU2 whi7-13AP-5FLAG::HisMX* |
| Kim 165 | *BY4741 WHI7-5FLAG::NatMX* |
| Kim 165 | *BY4741 whi7-13AP+3AQ-5FLAG::HisMX* |
| Kim 166 | *BY4741 rad53Δ::KanMX sml1Δ::Leu2 WHI7-5FLAG::NatMX* |
| Kim 188 | *BY4741 CKS1-13MYC::NatMX* |
| Kim 199 | *BY4741 cdc15-1 WHI7-5FLAG::NatMX* |
| Kim 209 | *BY4741 cdc15-1 whi7-13A-5FLAG::HIS3* |
| Kim 245 | *BY4741 whi7Δ::KanMX whi5Δ::HisMX SIC1-5FLAG::NatMX* |
| Kim 247 | *BY4741 CKS1-13MYC::NatMX WHI7-5FLAG::HIS3* |
| Kim 260 | *BY4741 mec1Δ::KanMX sml1Δ::LEU2 whi7Δ::URA3 CKS1-GBP::NatMX* |

| | |
|---|---|
| Kim 264 | *BY4741 SIC1-5FLAG::NatMX* |
| Kim 290 | *BY4741 mec1Δ::KanMX sml1Δ::LEU2 swi6Δ::NatMx* |
| Kim 291 | *BY4741 rad53Δ::KanMX sml1Δ::LEU2 swi6Δ::NatMx* |
| Kim 333 | *BY4741 SIC1-GFPmut1::kanMX* |
| Kim 335 | *BY4741 SIC1-GFPmut1::kanMX whi7Δ::URA3 whi5Δ:: HygR* |
| Kim 371 | *BY4741 GPDpr-3FLAG-MEC1::NatMX* |
| Kim 374 | *BY4741 WHI5-GFPmut1::NatMX* |
| Kim 376 | *BY4741 WHI5-GFPmut1::NatMX rad53Δ:: KanMX sml1Δ:: LEU2* |
| Kim 433 | *BY4741 whi7Δ::KanMX SWI6-FLAG::HIS3 SWI4-FLAG::NatMX* |
| Kim 490 | *BY4741 mec1Δ::KanMX sml1Δ::LEU2 swi4Δ:: HygR* |
| Kim 492 | *BY4741 rad53Δ::KanMX sml1Δ::LEU2 swi4Δ:: NatMX* |
| Kim 519 | *BY4741 CKS1-13MYC::NatMX whi7ΔC-5FLAG::HIS3* |

**Table S2. Plasmids used in this study**

| Plasmid | Base plasmid/Genotype |
|---|---|
| pRS426-*RNR3pr-WHI7* | *amp$^r$/URA3 RNR3pr-WHI7* |
| pRS426-*RNR3pr-whi7 WIQ* | *amp$^r$/URA3 RNR3pr- whi7 R128W A132I K135Q* |
| pRS426-*RNR3pr-WHI5* | *amp$^r$/URA3 RNR3pr-WHI5* |
| pRS426-*RNR3pr-whi5 WIQ* | *amp$^r$/URA3 RNR3pr- whi5 R185W A189I K192Q* |
| pRS426-GPD*pr-WHI5* | *amp$^r$/URA3 GPDpr-WHI5* |
| pRS426-GPD*pr-WHI7* | *amp$^r$/URA3 GPDpr-WHI7* |
| pRS426-GPD*pr-whi7 WIQ* | *amp$^r$/URA3 GPDpr- whi7 R128W A132I K135Q* |
| pGEX6p-1 *whi7(1-216)* | *amp$^r$ GST- whi7 1-216aa* |
| pRS313-p-WHI7 | *amp$^r$/HIS3 WHI7* |
| pRS313-p-*WHI7-GFP* | *amp$^r$/HIS3 WHI7-GFP* |
| pRS426-*RNR3pr-whi7(1-221)* | *amp$^r$/URA3 RNR3pr- whi7(1-221)* |
| pRS313-p-*whi7(1-216)* | *amp$^r$/HIS3 whi7(1-216)* |
| pET-15b-RAD53 | *amp$^r$ 6His-RAD53* |
| pET-15b-*rad53-KD(K227A)* | *amp$^r$ 6His-rad53-KD(K227A)* |
| pET-15b-RAD53 | *amp$^r$ 6His-RAD53* |
| pET-15b-*rad53-KD(K227A)* | *amp$^r$ 6His-rad53-KD(K227A)* |

| | |
|---|---|
| pGEX6p-1 WHI7 | *amp$^r$ GST-WHI7* |
| pGEX6p-1 Whi5 | *amp$^r$ GST-WHI5* |
| pGEX6p-1 *sic1*ΔC-*ALFA* | *amp$^r$ GST-sic1(1-216)-ALFA* |
| pGEX6p-1 *CKS1-ALFA* | *amp$^r$ CBP-CKS1-ALFA* |
| pRS426-*RNR3pr-whi7 3AQ* | *amp$^r$/URA3 RNR3pr- whi7 T222A,S224A,T225A* |
| pRS426-*RNR3pr-whi7 3DQ* | *amp$^r$/URA3 RNR3pr- whi7 T222D,S224D,T225D* |
| pGEX6p-1 *whi7 13D* | *amp$^r$ GST-whi7 T10D,S17D,S137D,S158D,S160D,S181D,S182D,S203D,S217D,S218D,T222D,S224D,T225D* |
| pRS426-*RNR3pr-whi7 13AP* | *amp$^r$/URA3 RNR3pr- whi7 T5,14,40,84,86,87,100A S27,56,62,98,105,212A-5FLAG* |
| pRS426-*RNR3pr-whi7 13D* | *amp$^r$/URA3 RNR3pr- whi7 T10D,S17D,S137D,S158D,S160D,S181D,S182D,S203D,S217D,S218D,T222D,S224D,T225D* |
| pRS426-*RNR3pr-whi7 13A* | *amp$^r$/URA3 RNR3pr- whi7 T10A,S17A,S137A,S158A,S160A,S181A,S182A,S203A,S217A,S218A,T222A,S224A,T225A* |

| | |
|---|---|
| pRS426-*RNR3pr-whi7 41A* | *amp$^r$/URA3 RNR3pr- whi7 alanine substitutions of all serine and threonine residues except 13 CDK sites(T5,T14,T40,T84,T86,T87,T100,S27,S56,S62,S98,S105,S212)* |
| pRS313-p-*whi7(1-216)-GFP* | *amp$^r$/HIS3 whi7(1-216)-GFP* |
| pRS426-GPD*pr whi7(1-216) WIQ* | *amp$^r$/URA3 GPDpr- whi7 R128W A132I K135Q* |
| pRS426-GPD*pr whi7 WIQ* | *amp$^r$/URA3 GPDpr- whi7 R128W A132I K135Q* |
| pRS316-p-4MYC-*whi7* | *amp$^r$/URA3 WHI7pr- whi7* |
| pRS316-p-4MYC-*whi7* | *amp$^r$/URA3 WHI7pr-4MYC-WHI7* |
| pRS316-p-4MYC-*whi7 WIQ* | *amp$^r$/URA3 GPDpr-4MYC-whi7 R128W A132I K135Q* |

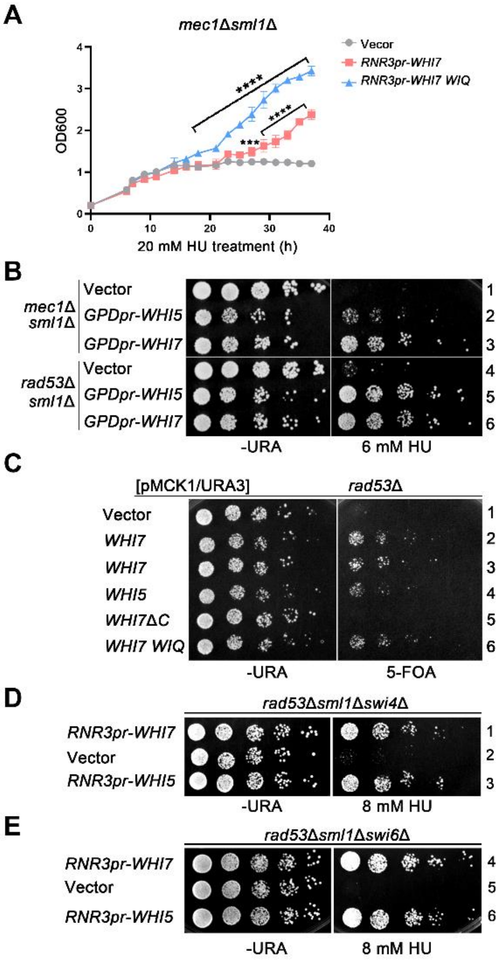


**Figure S1 (Related to Figure 1) Whi7/5 are positive effectors of Mec1-Rad53 checkpoint independently on G1/S transcription.**

(A) Overexpression of *WHI7* or *whi7*-WIQ suppresses the HU sensitivity of *mec1*Δ*sml1*Δ. The overnight cultures of the indicated strains were inoculated into fresh SC media (initial $OD_{600} = 0.2$) supplemented with 20 mM HU. $OD_{600}$ was measured every two hours from three independent repeats during the growth at 30°C. The statistical significance was calculated via two-way ANOVA analysis, ***, $p < 0.001$; ****, $p < 0.0001$.

(B) Constitutive overexpression *WHI7* or *WHI5* exhibits opposite effects on cell growth in the presence or absence of HU. The serial dilution assays were performed as described in Figure 1A, except that *WHI7/5* overexpression was driven by a constitutive GPD promoter.

(C) An extra copy of *whi7*-WIQ bypasses the essential role of *RAD53* as well. The indicated plasmid was transformed into the *mec1*Δ or *rad53*Δ haploid strain containing the pRS426-*MCK1* plasmid. Plasmid shuffling was conducted as described in Figure 1C.

(D, E) The checkpoint function of Whi7/5 is independent of either Swi4 (D) or Swi6 (E). *WHI7/5* was overexpressed in *rad53*Δ*sml1*Δ*swi4*Δ or *rad53*Δ*sml1*Δ*swi6*Δ. 5-fold serial dilutions were carried out as described in Figure1 G.

Kim_FigS2

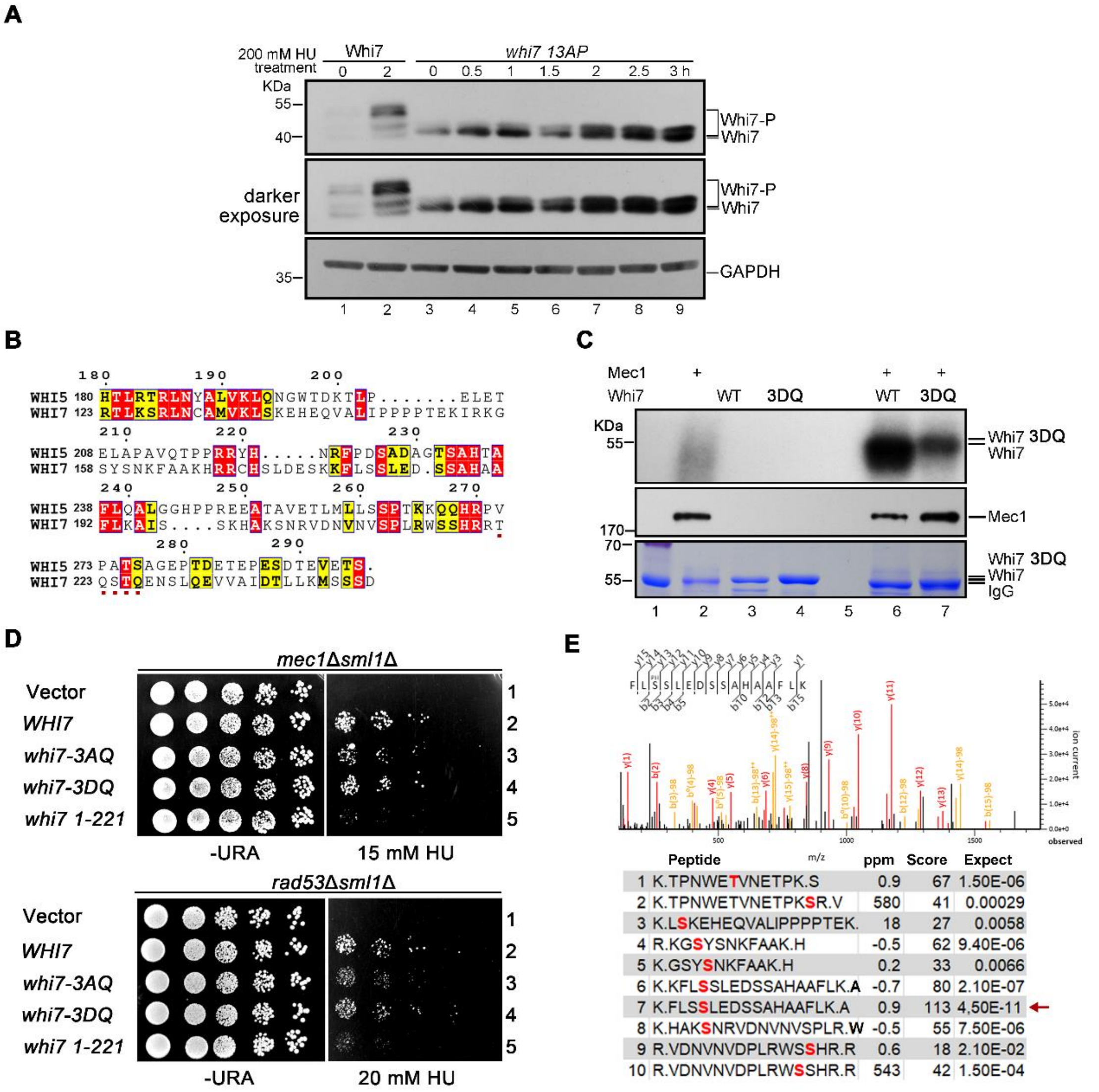


**Figure S2 (Related to Figure 2) Whi7 is phosphorylated by Mec1 and Rad53**

(A) HU induces Whi7 phosphorylation. WT, *whi7-13AP* cells were collected after 200mM HU treatment for the indicated time. The endogenous Whi7 carrying a 5FLAG tag in these cells was detected by immunoblots using an anti-FLAG antibody. whi7-13AP represents that a total of 13 potential CDK sites (S/TP) of Whi7 are substituted by alanine. GAPDH serves as a loading control.

(B) Whi7 contains an S/TQ cluster at the C-terminus. Whi7 and Whi5 protein sequences were aligned by CLUSTER W with default settings.

(C) Mec1 phosphorylates Whi7 *in vitro*. 3FLAG-Mec1 was precipitated from yeast cells and incubated with purified recombinant GST-Whi7 or GST-Whi7-13D in the presence of ɣ-$^{32}$P-ATP. In vitro kinase assays were conducted as described in Figure 2D. The 3FLAG-Mec1 band was confirmed by immunoblots using an anti-FLAG antibody.

(D) Phosphorylation of Whi7 by Mec1 contributes to dealing with replication stress. Various *whi7* phosphorylation mutants or truncations were overexpressed in *mec1*Δ *sml1*Δ or *rad53*Δ*sml1*Δ. 3AQ:T222A, S224A, T225A; 3DQ: T222D,S224D,T225D.

(E) The phosphorylation sites of Whi7 detected in MS/MS.

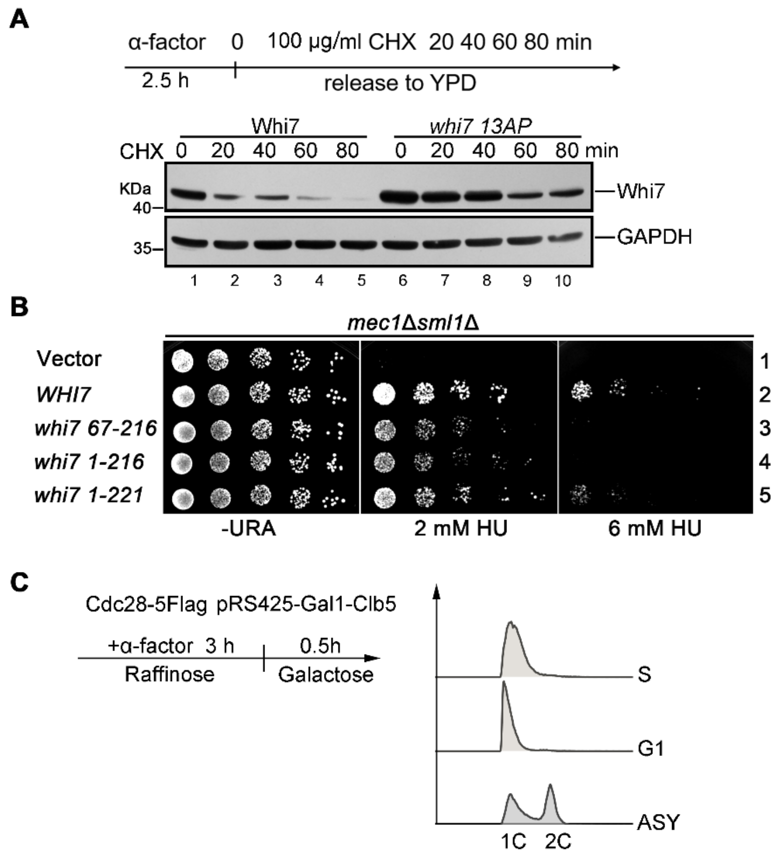


**Figure S3 (Related to Figure 3). The half-life of Whi7 is prolonged in response to replication stress**

(A) Whi7-13AP, no longer a substrate of CDK-dependent, was degraded more slowly than Whi7 WT during the normal cell cycle. Cells carrying the endogenous Whi7-5FLAG were arrested in G1 by α-factor and released into fresh media containing 0 or 200 mM HU. 100 μg/ml cycloheximide (CHX) was then added at t = 0; samples were taken every 20 mins. Cell lysates were analyzed by immunoblotting. GAPDH serves as a loading control.

(B) Various *whi7* truncation mutants were overexpressed in *mec1*Δ*sml1*Δ and subjected to 5-fold serial dilution assays.

(C) Clb5 was overexpressed under the control of the *GAL1* promoter in yeast cells bearing 5FLAG-tagged Cdc28 at its genomic locus. To further enrich S-CDKs, cells were released from α-factor for 30 min when most cells were in the early S phase, as evidenced by FACS.

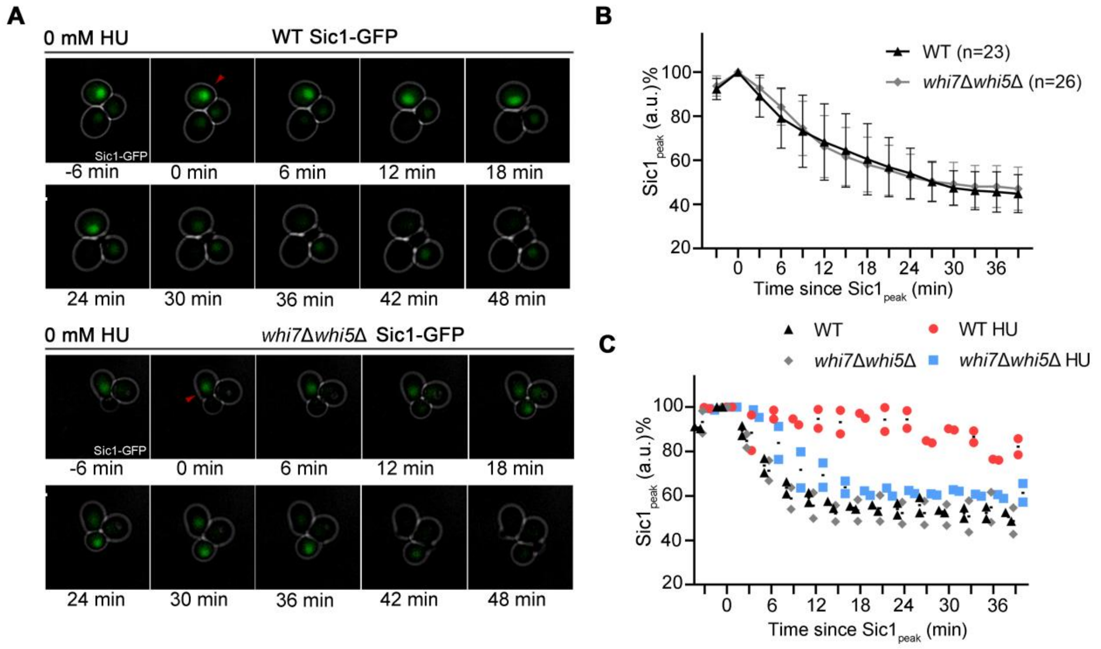


**Figure S4 (Related to Figure 5) Whi7 and Whi5 are involved in regulating Sic1 degradation in response to replication stress.**

(A, B) Single-cell live imaging of Sic1-GFP yeast cells without HU treatment.

(C) Single-cell profiles of Figure 5C and Figure S4A.

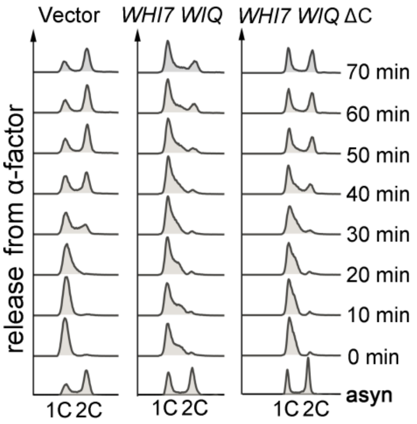


**Figure S5 (Related to Figure 5) Whi7 also regulates the normal cell cycle progression in a transcription-independent way via its C-terminus.** Cells were synchronized in G1 by α-factor before releasing into the fresh medium for the indicated times. The cell cycle progression was analyzed by FACS.
